# Coronavirus spherules are supported by a novel triple-layer protein lattice

**DOI:** 10.64898/2026.10.09.757900

**Authors:** Patrick M. Phillips, Joanna Wells, Naveen Mathew, Nicole Doyle, Sarah Keep, Parijat Majumder, James B. Gilchrist, Philippa C. Hawes, Jonathan M. Grimes, Maria Harkiolaki, Daniel K. Clare, Helena J. Maier

## Abstract

Coronaviruses (CoVs) cause significant disease in humans and animals, with a track-record of cross-species transmission. Despite their global significance, available control strategies are limited, especially for emerging CoVs. A deeper fundamental understanding of the virus life cycle is essential to underpin development of control measures. A critical step of the CoV life cycle is the rearrangement of host cell membranes into replication organelles (ROs). This is conserved across all CoVs and ROs comprise double membrane vesicles (DMVs), the site of viral RNA synthesis, and double membrane spherules. CoV spherules are complex structures requiring both positive and negative curvature of paired membranes so are predicted to have a high energy cost to maintain. To date they have an entirely unknown function. Interestingly, although always present, abundance of spherules in CoV infected cells varies with *Betacoronavirus* Middle East Respiratory Syndrome coronavirus, *Gammacoronavirus* infectious bronchitis virus (IBV) and porcine deltacoronavirus in particular producing a considerable number. Here, using cryo-bioimaging we identified for the first time that CoV spherules are embedded with a complex and organised triple-layered protein lattice and that spherules always contain a cargo that appears arranged as a shell inside the vesicle. In addition, we confirmed the presence of a double membrane spanning molecular pore in *Gammacoronavirus* DMVs and further identified a population of potential DMV formation intermediates. Together, this work provides a foundation for understanding a fundamental yet unresolved aspect of coronavirus replication.

## Introduction

Coronaviruses (CoVs) are a family of large single-stranded positive-sense RNA (+RNA) viruses. CoVs infect a broad range of species with members of the *Alpha-*and *BetaCoV* genera infecting mammals, including humans, and *Gamma-*and *DeltaCoV* genera infecting predominantly avian species. The economic and health impacts of CoV infections are extensive. Furthermore, due to their proven capability to jump between species, there is a continued threat of emergence of new CoVs into either humans or livestock with potentially devastating results on human and animal health, economic growth and food security globally. To underpin future development of pan-CoV control measures and improved preparedness for emerging CoVs, a detailed fundamental understanding of the conserved elements of the virus life cycle and virus-host cell interactions is essential.

All +RNA viruses, including CoVs, compartmentalise viral RNA synthesis into replication organelles (ROs), which are derived from modified host membranes^1,2^. The conserved elements of CoV ROs are double membrane vesicles (DMVs) and double membrane spherules tethered to either convoluted membranes (CM)^3–6^ or zippered ER (zER)^7–10^. DMVs have been a major focus of previous study. They are spherical vesicles with diameters of 250-300 nm^7,10,11^ and are known to be the primary site of RNA synthesis^6,7,10,12–14^. DMVs in some *BetaCoV* infected cells have now been shown to contain a double membrane-spanning protein pore, which provides a mechanism for RNA translocation between the DMV interior and the cytoplasm^15–17^. However, to date this pore has only been visualised in mouse hepatitis virus (MHV) and Severe Acute Respiratory Syndrome (SARS)-CoV-2 infected cells so the presence of the DMV pore in more diverse CoV remains unconfirmed.

In addition to DMVs, the conserved CoV RO also includes spherules^8,10,18^. Interestingly, although always present, spherule abundance varies between CoV species. While *BetaCoV* Middle East Respiratory Syndrome (MERS)-CoV, *GammaCoV* infectious bronchitis virus (IBV) and *DeltaCoV* porcine deltacoronavirus (PDCoV) produce abundant spherules^10^, *AlphaCoVs* porcine epidemic diarrhoea virus (PEDV) and HCoV-229E and *BetaCoVs* MHV, SARS-CoV and SARS-CoV-2 produce relatively few of these structures^10,19^. Spherules have diameters of 60-80 nm, have a uniform spherical shape and are always tethered to CM or zER. Their formation requires both positive and negative curvature of two membranes as well as maintenance of a neck where the spherule joins the zER or CM. Therefore, it is predicted that there is a high energy cost of manipulating and holding the host cell membranes in spherule networks. This energy cost, combined with their presence in all CoV infected cells, led to the hypothesis that they perform some necessary function during virus replication. However, as there is no data suggesting any direct role in viral RNA synthesis^10^, what their function may be remains elusive. Alongside observations of uniformity of spherules and their necks, earlier work also showed that they contain an electron dense material lining the inner membrane^8^, indicating that a protein scaffold is likely to be associated with spherules. However, the identity of any proteins associated with spherules is unknown. A more detailed understanding of the structure of this hypothesised scaffold may shed light onto the function of these conserved structures.

Here, we performed a comprehensive cryo-bioimaging analysis of the IBV RO. This demonstrated for the first time the significant extent of the RO at the late stage of infection, comprising hundreds of DMVs and spherules. We confirm the presence of molecular pores in *GammaCoV* DMVs, opening potential for future detailed structural comparison of this critical feature between different CoVs. Finally, cryo-electron tomography (ET) and subtomogram averaging (STA) has revealed a previously undocumented triple-layered protein lattice, including a cargo, associated with CoV spherules. Together this extends our understanding of the complexity of the CoV RO and will underpin further analysis of this critical and conserved step in the CoV life cycle, including future potential to manipulate these structures for viral control.

## Results

### Fluorescent recombinant IBV labels sites of nascent viral RNA synthesis

To directly visualise ROs, fluorescently tagged recombinant IBV (rIBV) with monomeric eGFP fused to nsp2^20,21^ was generated using reverse genetics^22^ and termed Beau-R-eGFP-nsp2 (Fig. 1a). Viral growth kinetics were measured in primary chick kidney (CK) and DF1 cells. Consistent with published data, WT virus titre peaked at 24 hours post infection (hpi) in CK cells (Fig. 1b(i)) and 48 hpi in DF1 cells (Fig. 1b(ii)). Beau-R-eGFP-nsp2 replicated to lower levels with peak titres 1log lower in CK cells (Fig. 1b(i)) and 2log lower in DF1 cells (Fig. 1b(ii)). However, replication dynamics were similar with peak titres reached at the same timepoint as WT virus. Therefore, although Beau-R-eGFP-nsp2 can replicate, insertion of the eGFP tag attenuates replication. To determine whether viral RNA synthesis was likely to be impacted by tag insertion, total levels of genomic RNA over a single replication cycle in CK cells were measured using qRT-PCR. Viral genomic RNA levels were not significantly different up to 12 hpi (Fig. 1b(iii)) indicating that the activity of the replication transcription complex (RTC) is unlikely to be impacted by the presence of the tag. Subsequently, to confirm that eGFP-nsp2 signal could be used as a marker to directly visualise replication organelles, eGFP colocalisation with nascent viral RNA metabolically labelled with 5-bromouridine (BrU) was assessed^13^. Significantly, eGFP-nsp2 signal showed good association and direct colocalisation with BrU-labelled viral RNA at 6 hpi (Fig. 1c). Further analysis of earlier timepoints using super-resolution STED microscopy demonstrated that eGFP-nsp2 puncta could be detected as early as 2 hpi with strong association with BrU labelled viral RNA from 3 hpi through to 6 hpi (Fig. 1d). Although eGFP-nsp2 and BrU RNA puncta were not directly colocalised, this is consistent with previous data^13^ and is consistent with nsp2 being localised on the cytoplasmic side of the DMV^23^.

**Figure 1:**
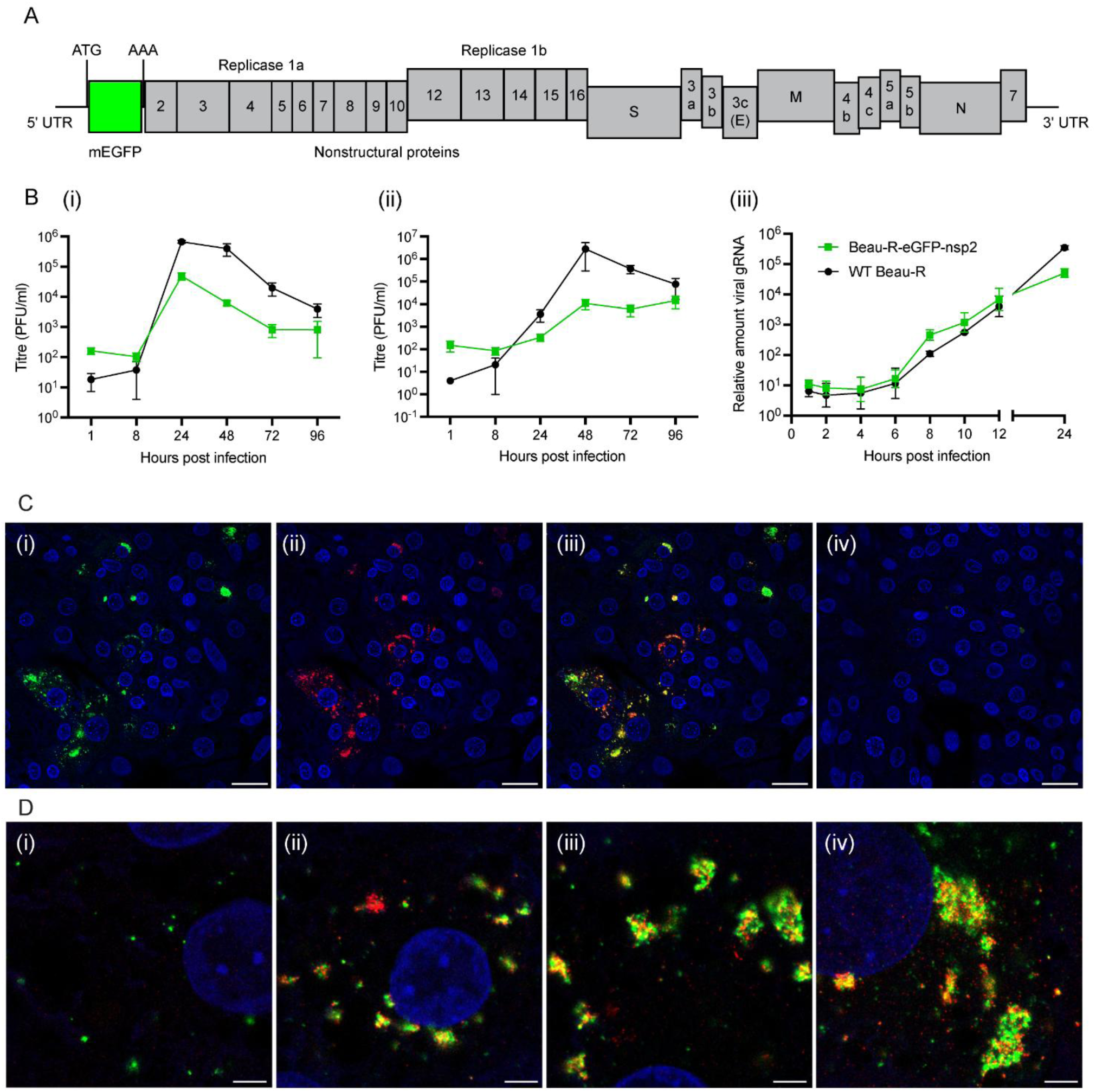
Beau-R-eGFP-nsp2 replicates and eGFP-nsp2 colocalises with sites of viral RNA synthesis. (**a**) Schematic of the genome of Beau-R-eGFP-nsp2, indicating the position of the meGFP insertion. (**b**) CK cells (**i**) or DF1 cells (**ii**) were infected with WT Beau-R (black) or Beau-R-eGFP-nsp2 (green). Progeny virus was collected at the indicated times and titrated by plaque assay. (**iii**) CK cells were infected with WT Beau-R (black) or Beau-R-eGFP-nsp2 (green) and total cell RNA was collected at the indicated times. Levels of viral genomic RNA (gRNA) were measured by qRT-PCR. Mean and SEM of three independent experiments are shown. (**c**) CK cells were infected (**i-iii**) with Beau-R-eGFP-nsp2 (green) or mock-infected (**iv**). Nascent viral RNA was labelled with BrU and cells were fixed at 6 h pi, labelled with anti-BrU antibody (red), and nuclei were stained with DAPI (blue). Scale bars indicate 25 µm. (**d**) CK cells were infected with Beau-R-eGFP-nsp2 (green) and nascent viral RNA labelled with BrU. Cells were fixed at 3 (**i**), 4 (**ii**), 5 (**iii**), and 6 (**iv**) h pi, labelled with anti-BrU antibody (red), nuclei were stained with Topro3 (blue), and imaged using the Leica SP8 gSTED microscope. Scale bars indicate 3 µm.

### Fluorescent-nsp2 localises to RO membranes

To observe whether eGFP-nsp2 localised to ROs and to confirm that RO appearance was not impacted by the presence of the tag, correlative light and electron microscopy (CLEM) with resin-embedded samples was completed. Fluorescence signal in infected DF1 cells at 18 hpi was shown to localise to ROs including DMVs, zER, and spherules (Fig. 2).

**Figure 2:**
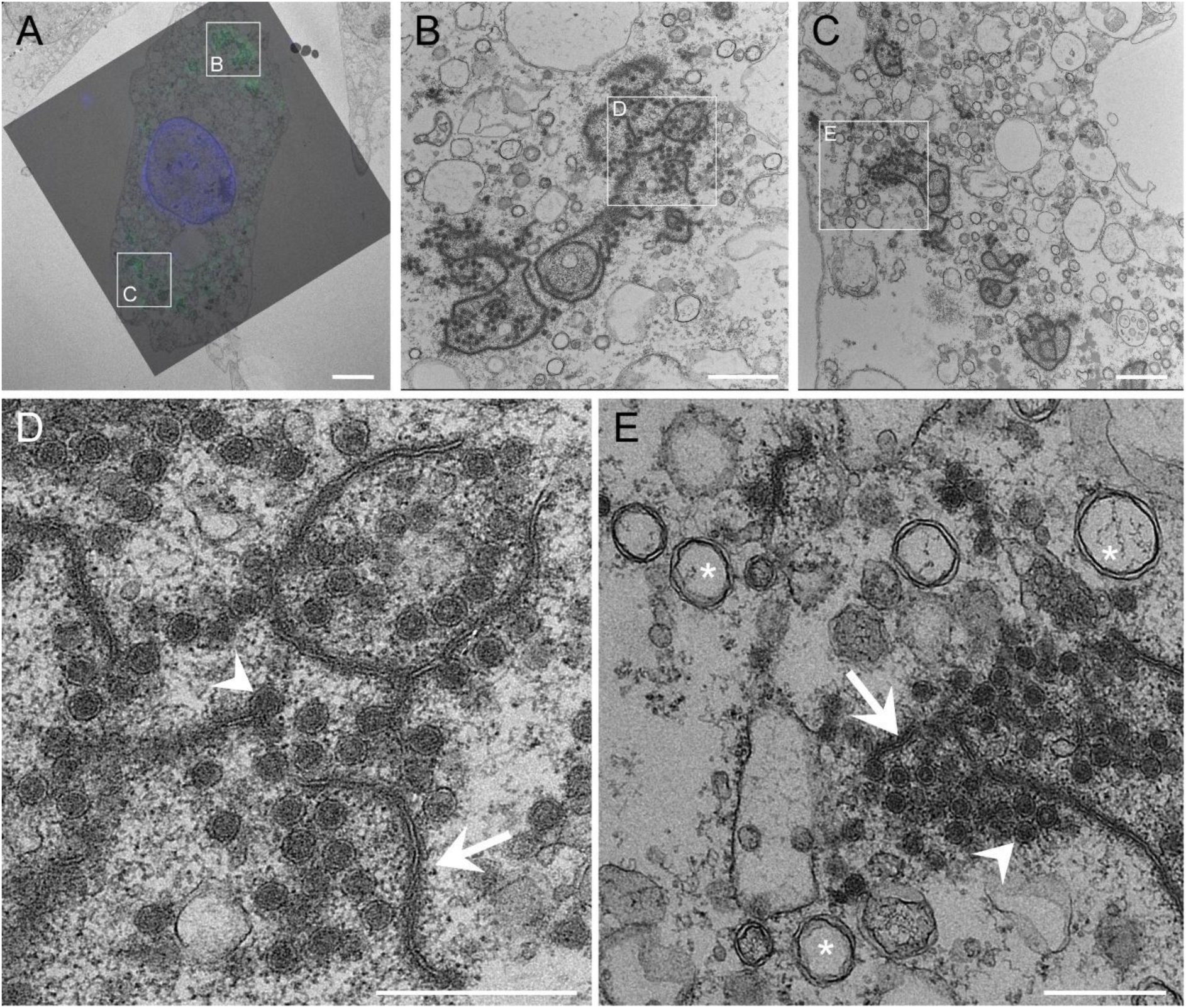
EGFP-nsp2 localises to IBV replication organelles. (**a**) DF1 cells were infected with Beau-R-eGFP-nsp2 (green) and nuclei were labelled with Hoechst (blue). Live cell fluorescence images were collected at 18 hpi and cells were then fixed and resin embedded. Infected cells identified during live cell fluorescence imaging were imaged by transmission electron microscopy. Fluorescence and electron microscopy images were overlaid. Higher magnification electron micrographs containing RO membranes (**b**, **c**) and (**d**, **e**) are shown. Note: (**b**, **c**) a different section through the same cell imaged in (**a**), at a different Z height, to more clearly show the accumulation of zER and spherules. Asterisks indicate DMVs, arrows indicate zER and arrowhead indicate spherules. Scale bars indicate 5 µm (**a**), 1 µm (**b**, **c**) and 500 nm (**d**, **e**)

Importantly, these were indiscernible from WT IBV ROs^8^. To visualise earlier ROs, Beau-R-eGFP-nsp2 infected cells were analysed using CLEM at 4-6 hpi (Fig. s1). At 6 hpi, DMVs were the predominant RO structure detected (Fig. s1b), with a small number of spherules also present (Fig. s1c). DMVs appeared fully formed with no formation intermediates detected (Fig. s1b). Cells imaged at 4 hpi, the earliest timepoint imaged to date, also contained DMVs but spherules were not detected at this timepoint (Fig. s1d). Together, this confirmed that Beau-R-eGFP-nsp2 is an appropriate tool to localise ROs.

### The IBV RO consists of DMVs and discrete patches of spherules

Although many studies of the CoV RO have been performed using fluorescence or electron microscopy, these approaches do not allow confident assessment of the full scale of the RO in the whole cells due to a lack of resolution or sample thickness, respectively. Therefore, whole Beau-R-eGFP-nsp2 infected cells were imaged using correlative fluorescence microscopy and cryo-soft X-ray tomography (SXT). At 24 hpi, Beau-R-eGFP-nsp2 fluorescence was efficiently detected and correlated to defined RO membranes imaged by cryo-SXT (Fig. 3). Spherical vesicles with diameters of 250 – 300 nm were imaged alongside darker sections of curved membrane with small (∼80 nm) dots tethered along the length (Fig. 3a). These structures likely correspond to DMVs and spherules, respectively and the darker appearance of the spherule-like membranes indicate they absorb more X-rays and have a higher carbon content. A single tomogram covered approximately one third of the total volume of the infected cell cytoplasm and 252 DMV-like vesicles and 424 spherule-like heads in 8 discrete patches of zER were counted. This cell also exhibited Beau-R-eGFP-nsp2 fluorescence and ROs in two further positions (Fig. 3a(i)), encompassing most of the cytoplasm so it could be estimated that over 750 DMV-like vesicles and over 1200 spherule-like heads existed in the cell. The average diameter of the DMV-like vesicles was 251 nm (n= 51, standard deviation = 27.9 nm) and the average diameter of the spherule-like heads was 72.4 nm (n= 56, standard deviation= 8.7 nm) (Fig. 3c). In another infected cell, a single eGFP-nsp2 fluorescence patch encompassed by one tomogram, contained 207 DMV-like vesicles and 372 spherule-like heads in 8 discrete patches. Overall, this demonstrates that by 24 hpi, the IBV RO is extensive and consisting of hundreds of DMVs that are interspersed with discrete patches of zippered ER (zER) with tethered spherules.

**Figure 3:**
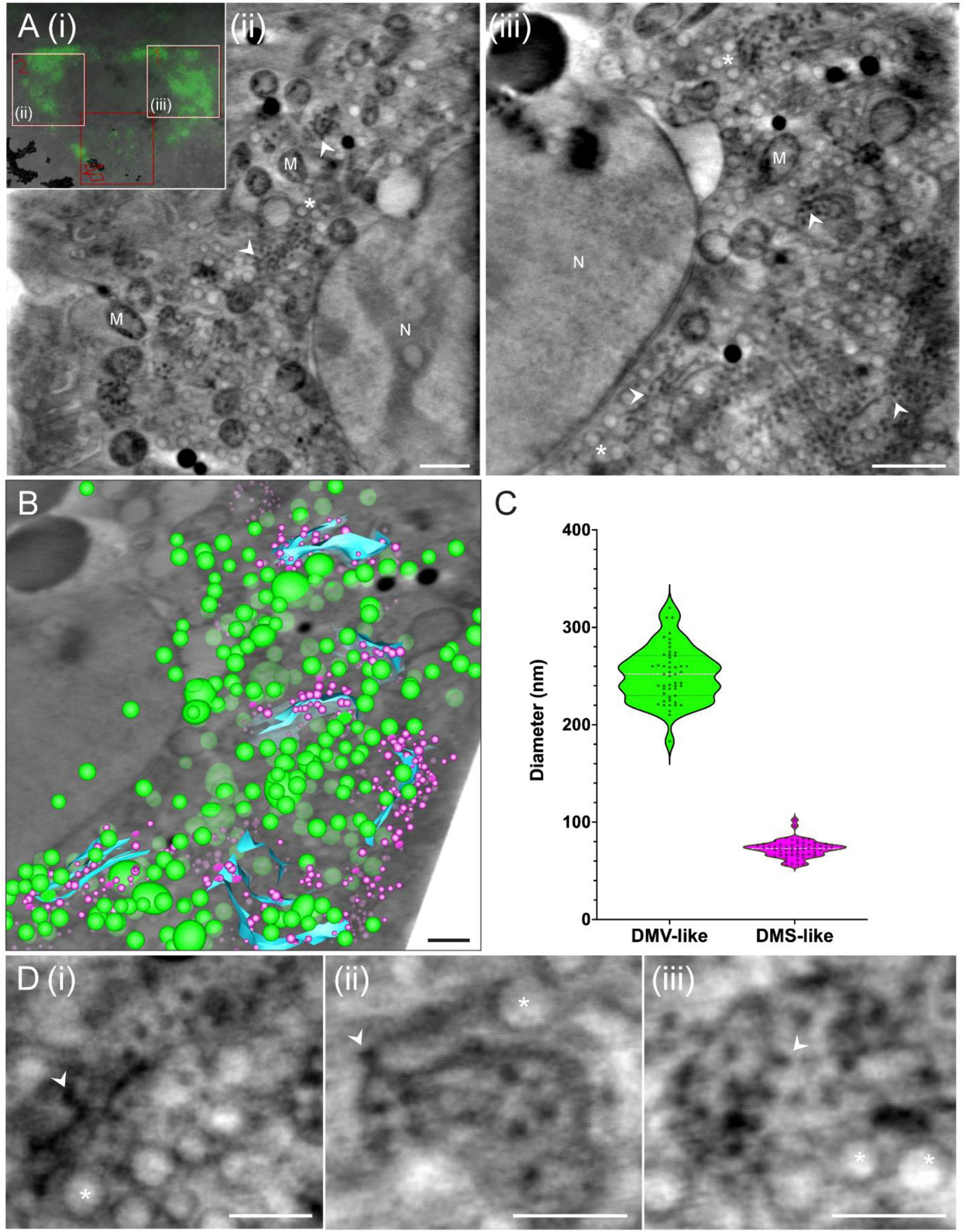
Cryo-soft X-ray tomography of Beau-R-eGFP-nsp2 infected cells. (**ai**) DF1 cells were infected with Beau-R-eGFP-nsp2 and plunge frozen at 24 h pi. Fluorescent regions observed via cryo-SIM were correlated to the same cell in the X-ray mosaic. Tomogram positions (**ii**) and (**iii**) were collected and reconstructed and a central orthoslice is shown for each tomogram. DMV-like membranes are marked with asterisks; spherule-like membranes are marked with arrowheads; M, mitochondria; N, nucleus. Scale bars indicate 10 µm (**i**) and 1 µm (**ii**, **iii**). (**b**) Segmentation of tomogram A(ii) with DMV-like membranes in green, spherule-like membranes in magenta, and sheets of putative zER labelled in cyan. Scale bar indicates 500 nm. (**c**) Diameters of DMV-like and spherule-like membranes from **b**. The median diameters are shown by horizontal white dashed lines, and the quartiles are shown by horizontal black dotted lines. (**d**) Gallery of DMV-(asterisks) and spherule-like (arrowheads) membranes taken from tomograms (**ii**) and (**iii**). Scale bars indicate 500 nm.

### IBV DMVs contain pores

DMVs have been imaged by cryo-ET in cells infected with *BetaCoV*s MHV and SARS-CoV-2 identifying a molecular pore spanning the double membrane^24^. However, there is no such data confirming the presence of a pore for other CoV genera, including *GammaCoV*s. DF1 cells infected with Beau-R-eGFP-nsp2 were vitrified at 24 hpi and eGFP-nsp2 fluorescent ROs were targeted for cryo-FIB/SEM to produce lamella for cryo-ET. In several reconstructed tomograms, spherules, zER, DMVs, and virus particles were observed (Fig. 4). Initial focus was on DMVs (Fig. 5) and consistent with our earlier work, the DMVs appeared as clusters of individual vesicles in the cytoplasm (Fig. 5a). Some DMV-ER connectors were observed and strand-like patterns consistent with RNA filaments were observed within the DMV interior. Significantly, for the first time for a *GammaCoV*, pores were observed spanning the DMV double membrane (Fig. 5). Pores appeared to be randomly and sparsely spread around the double membrane with just 1-3 pores per DMVs in the reconstructed volume. As whole DMVs were not included in the tomograms due to the thinness of the lamella and the missing wedge of tomographic data, it can be estimated that 5-10 pores exist per DMV. DMV pores comprised a visible membrane spanning region with cytosolic crowns and associated luminal and cytoplasmic densities, which are likely to be N, RNA, or other components of the RTC (Fig. 5d-g). The pores had a width of 20 nm within the membrane, and the cytosolic prongs extended 10 nm from the DMV outer membrane. Insufficient numbers of IBV DMV pores were visualised to allow STA in the current work so although their presence can be confirmed, a more detailed assessment of their structure was not possible.

**Figure 4:**
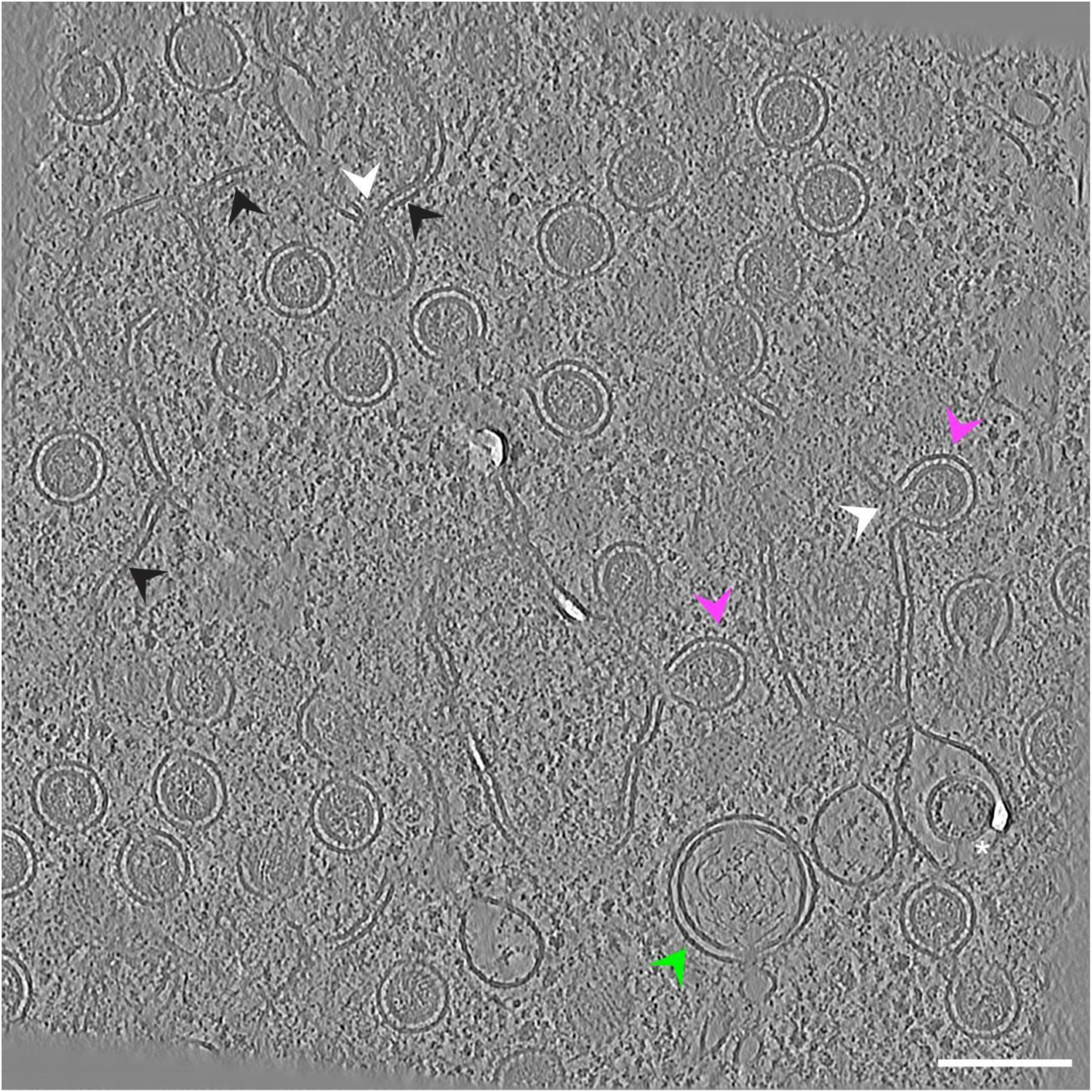
Cryo-electron tomography of the Beau-R-eGFP-nsp2 replication organelle. DF1 cells were infected with Beau-R-eGFP-nsp2 and plunge frozen at 24 h pi before correlative FIB milling to generate lamella for tomography. Sections of zippered ER (black arrowheads), spherule necks (white arrowheads), spherule heads (magenta arrowheads), single membrane vesicles containing virions (asterisk) and DMVs (green arrowhead) are indicated. Scale bar indicates 100 nm.

**Figure 5:**
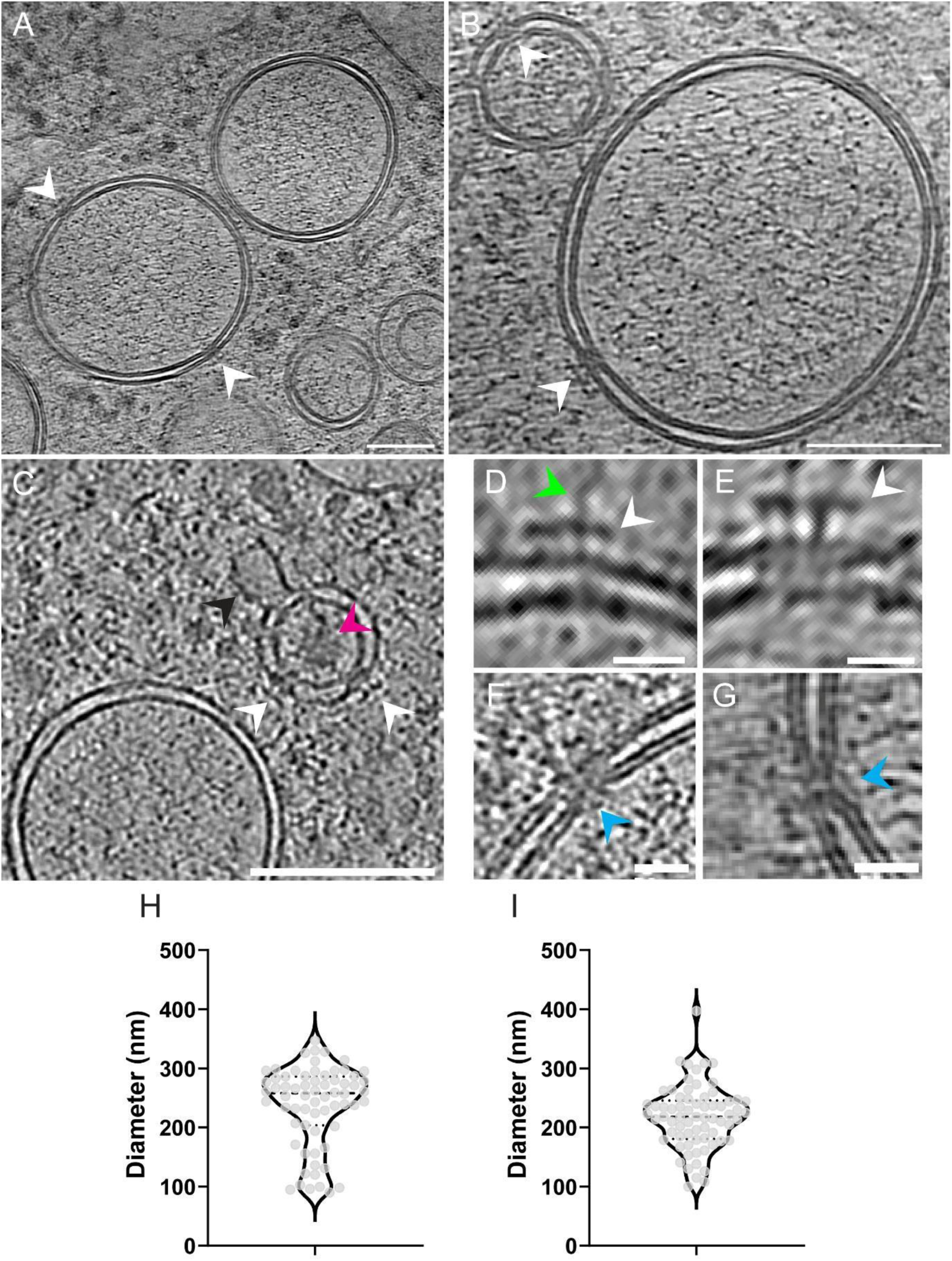
IBV DMVs contain double membrane spanning pores. (**a**) Beau-R-eGFP-nsp2 induced DMVs containing double membrane spanning pores (white arrowheads) and strand-like densities within their interior. Scale bar is 100 nm. (**b**) Small and large DMVs with double membrane spanning pores (white arrowheads). Scale bar is 100 nm. (**c**) Small DMV with pores (white arrowheads), attached single membrane sheet (black arrowhead) and dense content (magenta arrowhead). (**d-g**) Close up images of individual DMV pores showing crown-like structure on the cytoplasmic face (white arrowheads) and additional associated densities on the cytoplasmic (green arrowhead) and luminal (blue arrowheads) faces. Scale bars indicate 100 nm (**a**, **b, c**), 20 nm (**d**, **e**). (**h**) Maximum DMV diameters within the cryo-ET datasets were measured and plotted with mean (dashed line) and upper and lower quartiles (dotted lines) indicated. (**i**) CK cells were infected with WT Beau-R and fixed and resin embedded at 24 hpi. Cells were imaged using room temperature electron tomography.

### A population of small DMVs is present

The diameters of DMVs observed in the cryo-ET dataset was measured and DMVs had a mean diameter of 265 nm (± 36.3 nm, n= 53, Fig. 5h). The largest measured diameter was 347 nm and the smallest was 171 nm. Interestingly, a second population of small DMVs also containing double membrane spanning pores were observed alongside the larger DMVs.

Thirteen examples of small DMVs with diameters ranging from 90-166 nm were observed. These were of interest because they are comparable in size to minimal DMVs induced by the co-expression of *BetaCoV* nsp3-4 alone in the absence of other viral components^16,25^. These small DMVs had less uniformly spaced membranes than the larger DMVs but, significantly, a comparable number of visible pores. Moreover, the small DMVs contained some tightly packed content in their centres that appeared different to the overlapping RNA filaments observed in the larger DMVs (Fig. 5c). One example of a small DMV also appeared with a single membrane connector (Fig. 5c). To gain more information about the populations of DMVs with different sizes, CK cells were infected with WT Beau-R and processed for room temperature (RT) resin-embedded ET. Here, DMVs had a mean diameter of 217 nm (± 53.5 nm, n= 69) with the largest diameter measuring 397 nm and smallest measuring 100 nm (Fig. 5i). Therefore, both room temperature ET and cryo-ET data support that a range of sizes of DMVs exist, including a population of smaller DMVs.

### Spherule membranes are embedded with a protein scaffold

Subsequent focus moved to spherules within the cryo-ET data. Due to their complex structure requiring both positive and negative curvature of paired membranes to form the spherule body and neck, as well as earlier observations of electron dense material lining the inner membrane and higher X-ray absorption compared to DMV membranes, it seemed probable that a protein scaffold is associated with spherule membranes. Within tomograms, sections of zER were observed to run into spherule necks that curved into spherule heads (Fig. 4, black, white and magenta arrowheads). The spherules were uniform in size, always appeared with material in their interiors, and were always tethered to zER. Some longer regions of zER appeared without spherules while large bunches of several spherules tethered to zER were also observed. These observations were all consistent with those from cryo-SXT and previous room temperature EM and ET of IBV ROs^8,26^. Strikingly, a previously unreported striation pattern of electron density spanning the zER and spherule double membranes was observed, here referred to as spokes (Fig. 6). Both individual and clusters of DMVs were also observed in proximity with spherules, however, they did not contain any striation pattern between the membranes, demonstrating that this is a spherule specific structure and further indicating that the DMVs and spherules are derived via different mechanisms.

**Figure 6:**
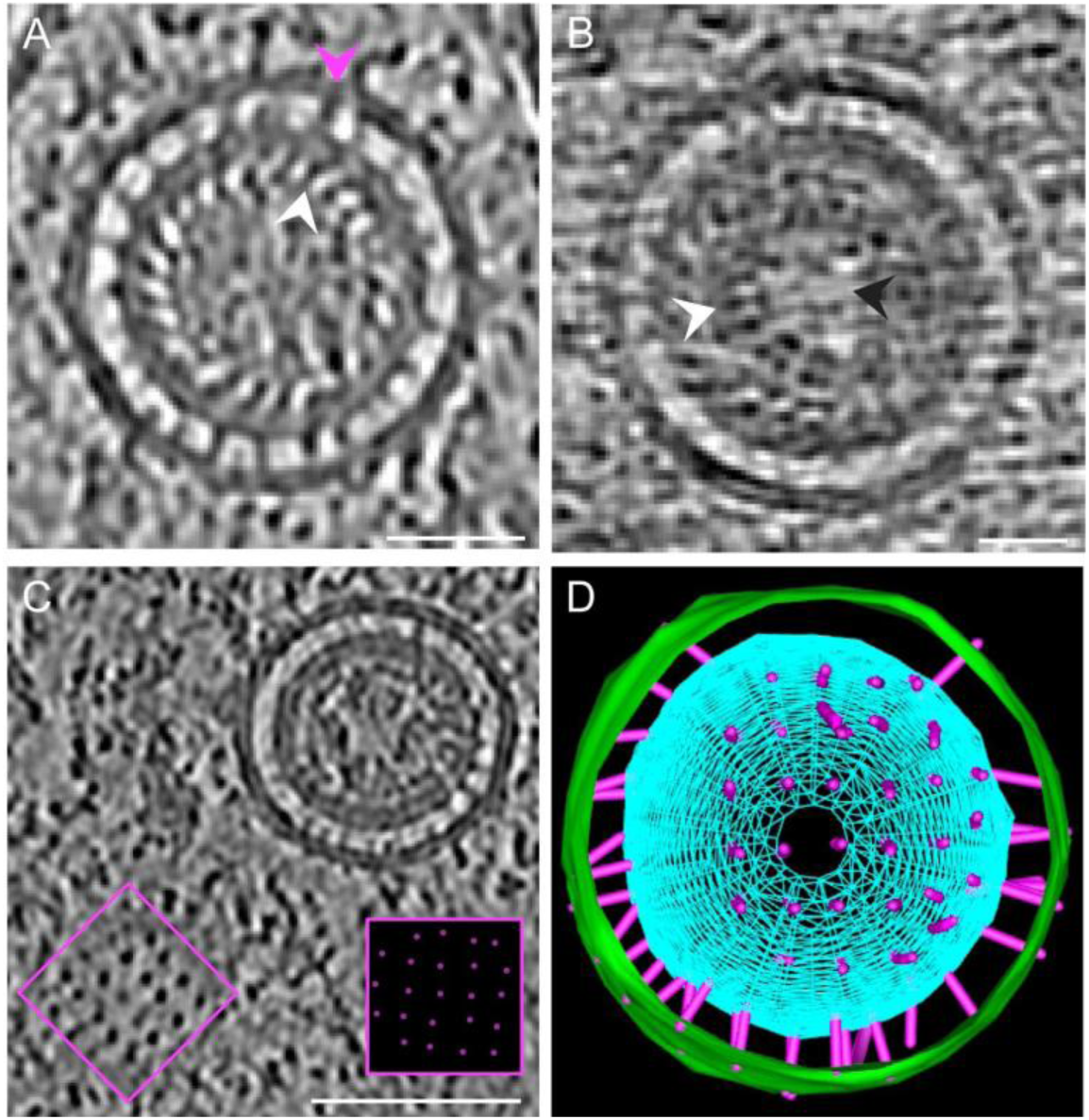
Spherules are embedded with a scaffold of intermembrane spokes and associated z-layer and cargo lining the inner membrane. (**a**) Spherule cross-section with evenly distributed spokes (magenta arrowhead) between the double membrane and ordered striation patterns laying at an angle lining the spherule inner membrane, which we have named the ‘z-layer’ due to the orientation of the densities (white arrowhead). (**b**) Spherule cross-section with cargo (white arrowhead) and lower electron density in the spherule centre (black arrowhead). (**c**) Axial view of a spherule showing the inter-membrane space (boxed) revealing the spokes in an ordered lattice (spoke arrangement highlighted in magenta in cut away). (**d**) IMOD segmentation of a spherule with the spokes (magenta), the outer membrane (green), and the inner membrane (cyan) which has been extrapolated into a sphere shown as a mesh. Scale bars indicate 20 nm (**a**, **b**) and 50 nm (**c**).

The spokes spanning the spherule double membrane appeared to extend into the spherule interior forming a second layer of striations under the inner membrane at an angle to the spokes, termed the z-layer (Fig. 6a). Below the z-layer, was a third layer of density, termed the cargo layer, with a degree of repeating structure and appearing as a shell with a gap in density at the spherule centre (Fig. 6b). Although there were some additional densities on the cytoplasmic surface of the spherules, they did not appear continuous with the spokes. The spokes spanning the spherule double membrane appeared to form ordered grid-like patterns as shown in the segmentation of the axial view of the intermembrane space (Fig. 6c, d) with an average distance between adjacent spokes of 6 nm (n= 52). Due to the thinness of the lamella and the missing wedge of tomographic data, spokes could not be resolved around the full spherular double membrane. However, an average of 68.7 (n= 10) spokes were visible covering one sixth of the surface area of the full spherular double membrane. Therefore, assuming that the spokes were equally spaced around the whole spherule head then it could be estimated that approximately 400 spokes exist in each spherule.

### Spherule necks can be open or sealed but lack a cap or crown

Focussing on the spherule necks, most of necks appeared open but there were a range of morphologies including previously undocumented longer or extended necks (Fig. 7a-c). Of the spherules with visible necks (Fig. 7a(i)), the average neck width was 21 nm (± 3.5 nm) (n= 61). However, seven examples of spherules with thinner necks were imaged where the spherule inner membrane appeared to be closed while the outer membrane was continuous with the zER (Fig. 7b(i)), although it cannot be ruled out that these necks could have an opening outside of the reconstructed volume of the tomogram. Finally, in the examples of the spherules with extended necks, the outer membrane appeared tightly apposed along the length of the extended neck (Fig. 7c) with a sealed inner membrane around the spherule head. Therefore, 90% (n= 68) of the visible necks were open and 10% of the spherules were or may be sealed. Interestingly, at this resolution, there were no obvious caps or crowns holding open or stabilising the open necks.

**Figure 7:**
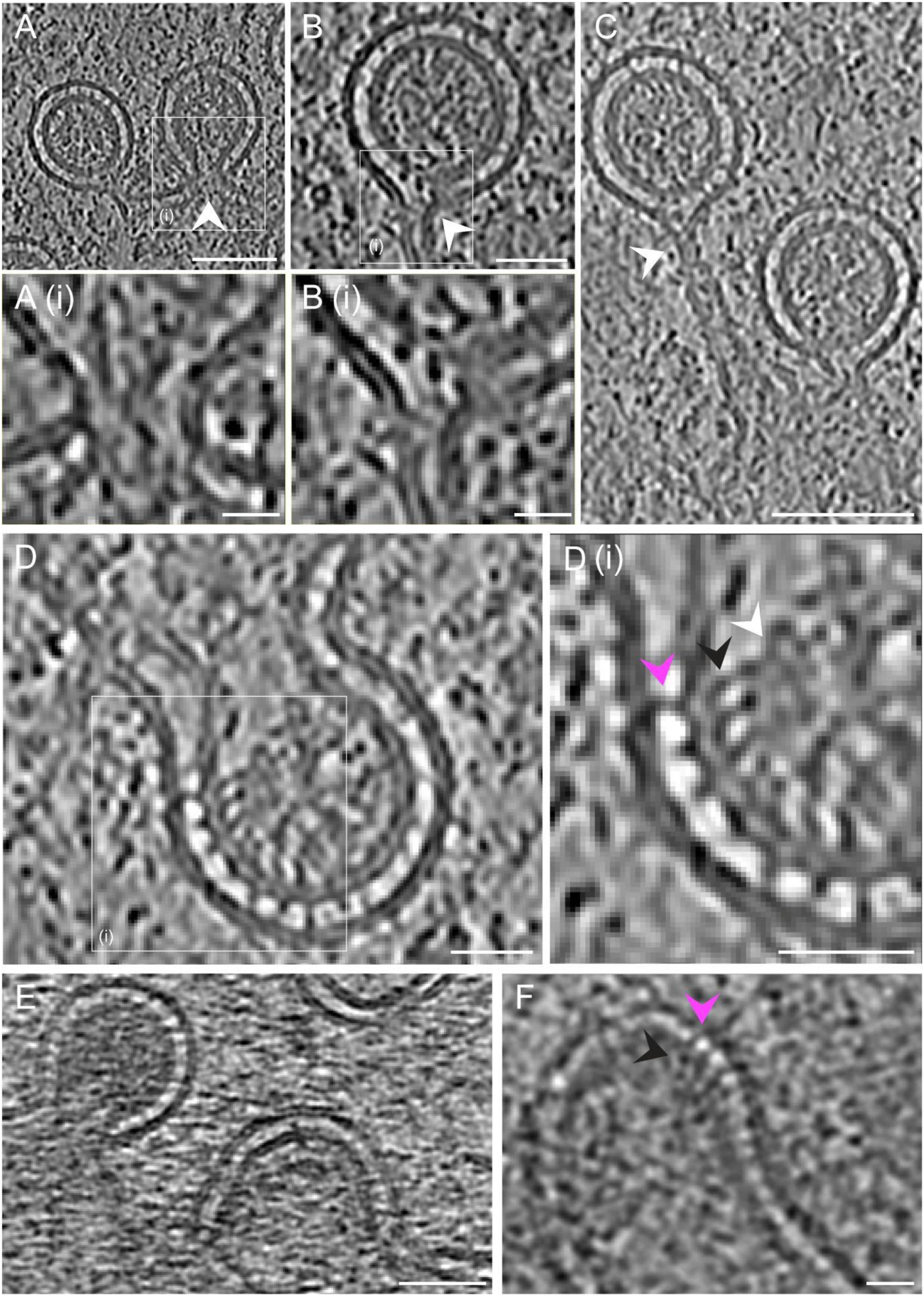
Spherules have open or sealed necks and contain cargo. Spherules with open (**a**, white arrowhead) or sealed (**b**, white arrowhead) necks with expanded boxed areas (**ai**, **bi**). Spherule with long neck (**c**, white arrowhead). (**d**) Spherule with a wide neck and expanded boxed area (**di**) with visible spokes (magenta arrowhead), a second interior z-layer (black arrowhead), and a third interior cargo density (white arrowhead). (**e**) Spherules adjacent to curved zER. (**f**) Curved zER containing spokes (magenta arrowhead) and z-layer (black arrowhead). Scales bars indicate 20 nm (**a**, **b**, **d**, **f**) and 50 nm (**c**, **e**).

Three examples of spherules with wider necks were also imaged, which may represent spherule formation intermediates (Fig. 7d-f). Interestingly, the z-layer densities were not observed either side of the spokes in the zER (Fig. 4, black arrowheads) indicating that the densities lining the spherule inner membrane must be incorporated during the process of spherule formation. Consistent with this model, an example of partially curved zER was observed where the z-layer and cargo layer appear to have begun to form under the concave membrane (Fig. 7f, black arrowhead). As all spherules were observed to contain the z-and cargo layers, it is possible therefore that the spherules may form via zER wrapping around a protein shell (Fig. 7d and 6b). Interestingly, although most adjacent spherules pinched out from a single section of zER in the same direction making a bunch, there were instances of spherules pinched out in opposite directions on the same section of zER. This suggests that the z-layer and the cargo layer can bind to either side of the zER depending on the orientation of the spokes within the zER.

### Spherules are embedded with a triple-layered protein lattice

To gain a clearer understanding of the structures associated with spherules, STA was performed (Fig. 8). Spherules were picked in 34 tomograms using a spherical picking strategy on each spherule head, using a total of 266 spherules. The spokes were spaced 60 Å apart so sub volumes were extracted every 40 Å around the surface of each sphere to oversample the surface of the spherule. After alignment and classification of the extracted sub volumes, STA of 3796 sub volumes (Fig. 8a) were refined to yield a density map at 23 Å resolution (Fig. 8b, d-f, Fig. s2). This revealed that spherules contain ordered and layered lattices of protein. Moving from the exterior to the interior of the spherule, side views of the model show the outer membrane, the spokes with a length of 80 Å, the inner membrane lined with the z-layer extending another 80 Å into the spherule interior, and the cargo layer densities (Fig. 8b-c). In the intermembrane space, the base of the spokes at the inner membrane appeared more electron dense than the region which contacts the inner leaflet of the outer membrane. The inner membrane appeared more distorted than the outer membrane and exhibited a ruffled pattern. Axial views of the model showed the spokes organised into a square lattice spaced by 60-90 Å (Fig. 8d(i)), the z-layer proteins lining the inner membrane organised into an oblique lattice with a unit cell of 60 Å by 70 Å (Fig. 8e), and the cargo layer with densities spaced by 50 Å (Fig. 8e).

**Figure 8:**
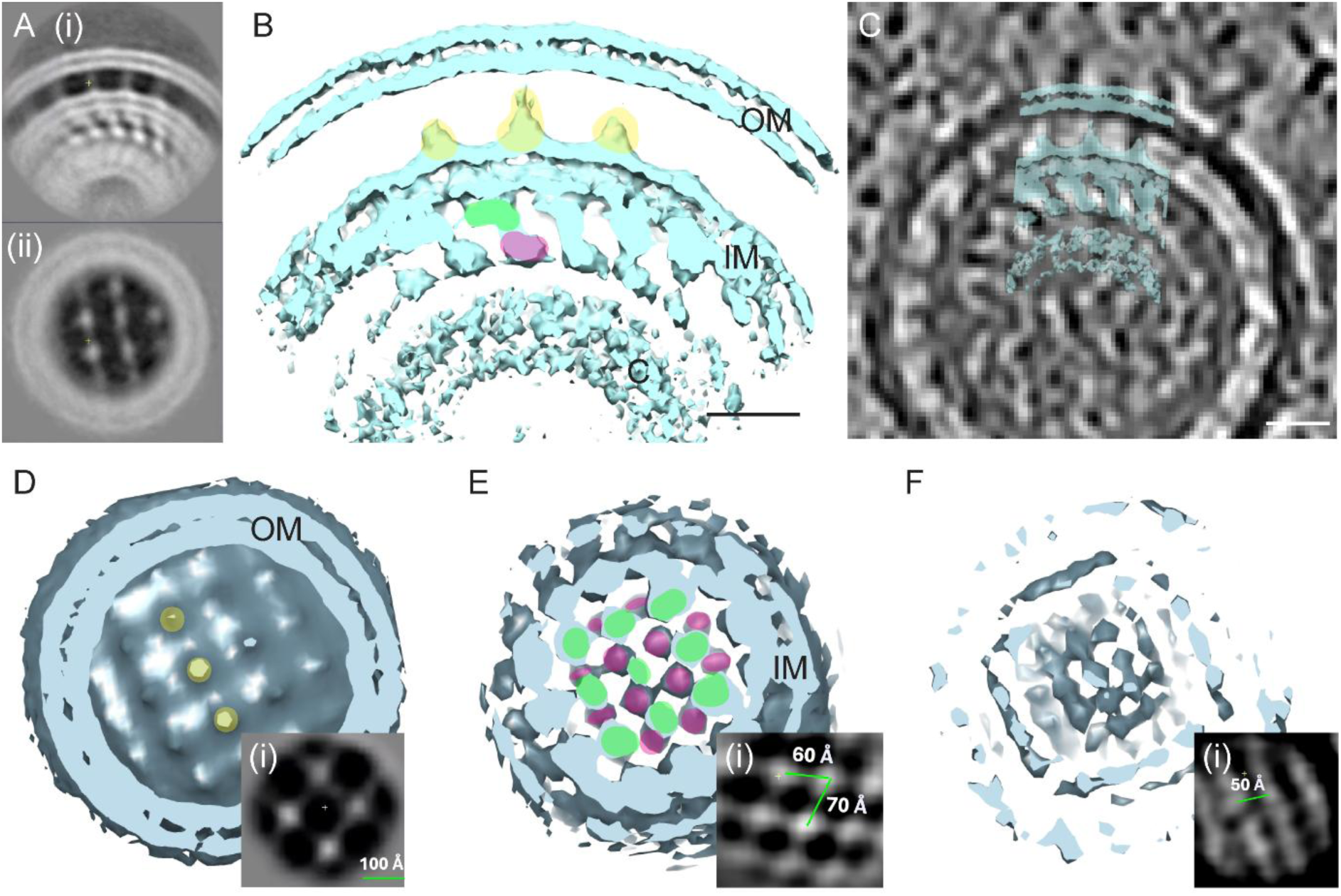
Subtomogram averaging reveals a triple-layer lattice supporting IBV spherules. (**a**) Side view (**i**) and axial view (**ii**) of the spherule particle average. (**b**) Average electron density of the side view of the spherule with three ordered layers: the outer membrane (OM), the spokes (yellow), the inner membrane (IM), the z-layer with upper (green) and lower (magenta) densities, and the cargo (C). Scale bar is 10 nm. (**c**) Overlay of the averaged spherule double membrane onto a representative spherule from the reconstructed tomogram. Scale bar is 10 nm. (**d**-**f**) Axial views of the STA model with the square grid pattern made by the spokes (**d**, yellow), the z-layer of densities labelled in green and magenta as in panel b (**e**), and the cargo (**f**). OM, outer membrane; IM, inner membrane. Insets (**di**, **ei**, **fi**) show the distances of the lattice-like densities in each layer.

### IBV structural proteins are unlikely to be constituents of spherule associated lattices

The identity of the proteins comprising the spherule associated lattice is not clear and the resolution achieved through STA does not allow prediction based on structure. However, based on spherule diameter being similar to that of a virus particle, it seemed possible that spherules may result from the expression of excess viral structural proteins. Therefore, to determine whether the spherule STA densities could be viral structural proteins, IF was performed. CK cells were infected with Beau-R-eGFP-nsp2, providing a marker for both DMVs and spherules. Cells were then labelled with antibodies targeting each of the IBV structural proteins, S, E, M, and N, and colocalisation with eGFP-nsp2 was analysed (Fig. 9). Using fluorescence intensity tracing over linear regions of infected cells, there was no consistent pairing of fluorescence peaks for eGFP and any of the 4 structural proteins, indicating limited to no colocalisation. Therefore, eGFP-nsp2 puncta were distinct from structural protein puncta and the structural proteins are unlikely to be localised to spherules or be constituents of the spherule associated lattice. Instead, it was hypothesised that viral nonstructural or accessory proteins or host cell proteins make up the spherule lattice. Due to a lack of antibodies targeting the other IBV proteins, a more detailed investigation to provide further insight into the identity of the spherule associated proteins was not possible.

**Figure 9:**
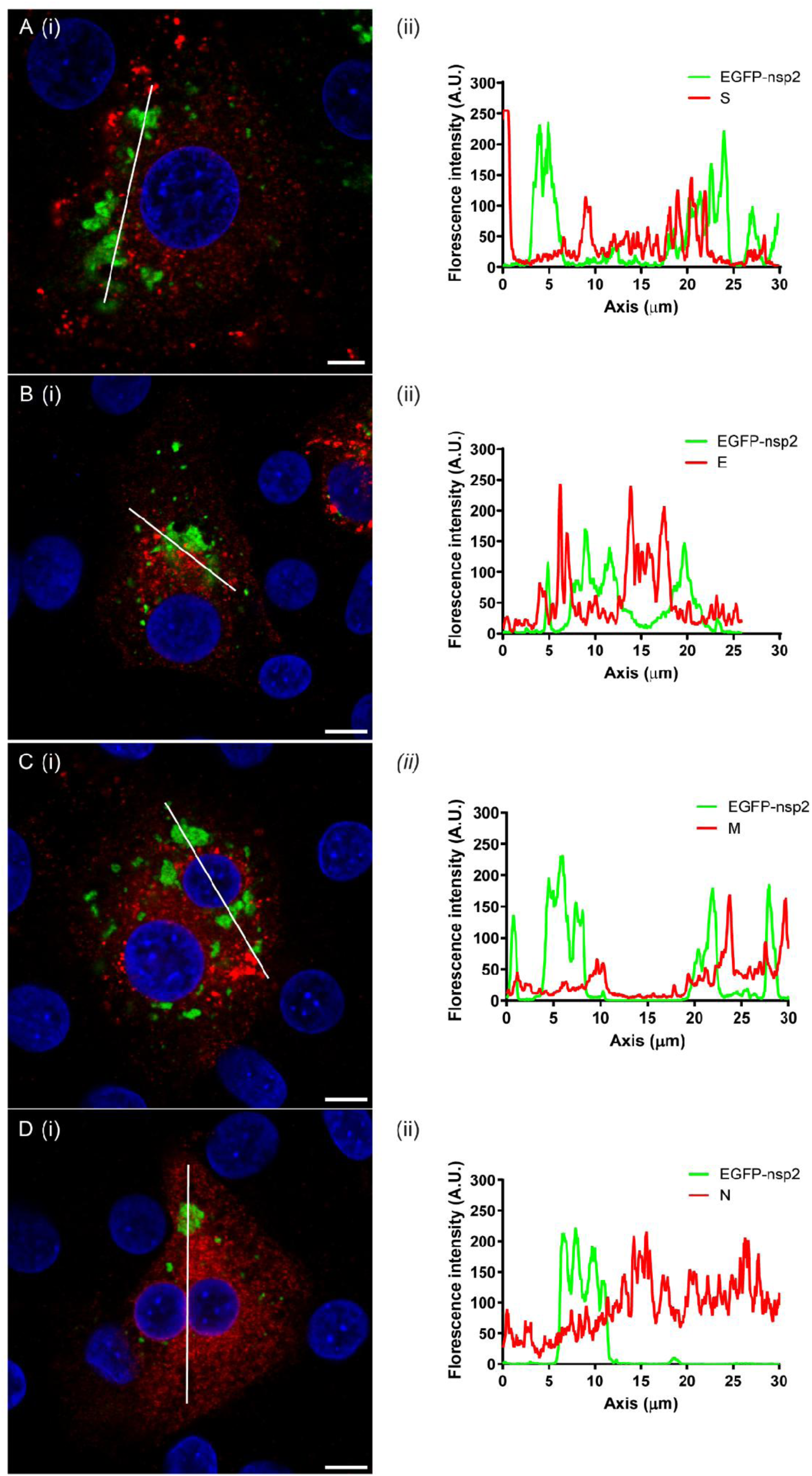
IBV structural proteins do not colocalise with ROs, including spherules. CK cells were infected with Beau-R-eGFP-nsp2 (green) and at 24 hpi were fixed and labelled to detect (**a**) spike (S, red), (**b**) envelope (E, red), (**c**) membrane (M, red) and (**d**) nucleocapsid (N, red). Nuclei were stained with DAPI (blue). To assess colocalisation of proteins, green and red fluorescence intensity along the indicated line in each panel (**i**) is plotted in panel (**ii**). Scale bars indicate 5 µm (**a**) and 8 µm (**b**-**d**).

## Discussion

Replication organelles are an essential and highly conserved feature of the CoV replication cycle. As such, they have been the focus of much study resulting in confirmation that DMVs provide the site for viral RNA synthesis with *BetaCoV* DMVs containing a molecular pore to allow RNA translocation and an increasing understanding of the organisation of DMVs and the viral and cellular proteins involved in their function and formation^16,23,25,27–30^. A less frequently studied element of the conserved CoV RO is spherules. In the current study, the structure of spherules has been revealed in unprecedented detail providing several novel insights. Our earlier RT EM of spherules suggested that they contain material and have more tightly apposed double membranes than DMVs^8,9,31^. This is consistent with cryo-SXT data generated here showing that the spherule-like membranes and zER absorb more X-rays than the DMV-like membranes, inferring that they have a higher carbon content and so are likely protein rich. Significantly advancing our understanding of spherules, cryo-ET revealed a striation pattern of spoke-like proteins spanning the spherule and zER double membranes. Subsequent STA revealed that the protein was ordered and formed a lattice inside spherules, which has not been observed for any other CoVs to date. This lattice was subdivided into three layers; a layer of spokes linking the two membranes of the spherule, the z-layer sitting under the inner membrane at a diagonal angle from the base of the spokes and the cargo layer making up a ring or shell in the interior of the spherule. Significantly, although spokes were visible linking the zER membranes, they were never seen on the DMV double membranes, consistent with earlier cryo-ET of *BetaCoV* DMVs^11^ and identifying the spokes as an entirely novel structure. The absence of spokes on DMV membranes demonstrates that DMVs form via a different mechanism to spherules. Interestingly, although IBV DMVs consistently appear as discrete vesicles with only rare external membrane connections, *BetaCoV*-induced CM (to which *BetaCoV* spherules are tethered) and DMVs form a continuous membrane network^5,7^. Therefore, it would be interesting to determine whether the spoke-like proteins seen here in IBV-induced zER exist in CM. This must be the case to allow spherule formation but there must also be spatial segregation to provide spoke-free membranes for DMV formation. In the current work, where the zER intermembrane space was resolved to a high enough level, all zER appeared to contain spokes (Fig. 4, black arrowheads) so it is unclear how ER membranes are targeted for either spherule or DMV formation. This so far unknown mechanism may provide an explanation for why some members of the CoV family form CM and others form zER. It is possible that higher expression or more concentrated insertion of spoke proteins into ER membranes results in longer tracks of zER (as in IBV infected cells), while lower expression or less organised insertion of spoke proteins into ER membranes would result in more reticulated CM (as in e.g. MHV infected cells). Further, this may also provide an explanation for the varying abundance of spherules between CoV species as more abundant spoke protein would presumably facilitate spherule formation.

Interestingly, there were no cap or crown structures resolved at the spherule necks. We had earlier hypothesised that to maintain the neck shape with tight curvature of paired membranes, that a protein complex would sit on the neck, like those observed for CHIKV and FHV single membrane spherules^32–34^. However, spherules produced by those viruses act as the site of viral RNA synthesis and the crown is comprised of the viral RNA synthesis machinery. Earlier work demonstrated that CoV spherules do not play a direct role in RNA synthesis^31^. In agreement, in the current study RNA filaments were never observed to be associated with spherules, unlike the DMVs. This emphasises that the CoV spherules are distinct from other +RNA virus spherules with a distinct function that is not RNA synthesis. Therefore, it is perhaps not surprising that there was no CHIKV or FHV-like crown on the IBV spherule necks. Instead, the spokes that were resolved between the spherule double membrane appeared to wrap around a uniform protein shell forming the cargo of the spherule. If this interaction is sufficiently stable, this could explain how the shape of the membrane is maintained without a cap sitting on the neck. Based on the observation that spherules always contained a cargo and were never empty, here we hypothesise a model for spherule formation in a stepwise fashion: 1) two ER leaflets are zippered by the spoke-like proteins, 2) another protein or complex of proteins interacts on one face of the zER and triggers oligomerisation of a uniform lattice making the z-layer and cargo of the spherule. In this model, the zER wraps around the cargo analogous to a clathrin or COPII coated vesicle but on the inside rather than the outside of a budding membrane.

Although the identity of the protein(s) forming zER and spherules remains unknown and their function remains elusive, it is possible to hypothesise which proteins may make up the spherule lattice. STA of virions and virus-like particles have described S, the RNP, and M producing layered lattices like those observed in spherules here^35–37^. However, immunofluorescence imaging for these structural proteins suggested that they are unlikely to localise to ROs. As DMV formation is known to involve nsp3, nsp4 and nsp6^25,27,29,30^, these nsps are likely to be involved in formation of other elements of the RO but in potentially different conformations or complexes. Nsp4 was thought to be a likely candidate for the spokes because IBV nsp4 alone is known to zipper ER membranes^38^. Interestingly, cryo-ET structures of nsp4 within DMV pores have shown it spanning the double membrane of the DMV^16,28^. Based on this structure and an AlphaFold predicted structure of IBV nsp4 it is possible to see how nsp4 would potentially be able to traverse both membranes of the spherule with an appropriately sized ectodomain and the nsp4 CTD contributing to density lining the inner membrane of the spherule (Fig. s3). However, a higher resolution EM density map from further cryo-ET and STA studies in the future will be required to fully determine the structure of the spherule spokes. The identity of the z-layer and cargo layer of the lattice are more difficult to hypothesise at this time. It is known that many of the nsps from *BetaCoV*s localise to the CM in addition to the DMVs, so these are possible candidates^3,5,7,31^. It is also possible that viral accessory proteins or even cellular proteins localise to spherules. Overall, identification of these proteins in the future will be valuable to give clues to the function of spherules and will likely provide insight into why abundance of spherules varies between CoV species.

In addition to the novel spherule associated lattice, cryo-ET revealed molecular pores spanning the IBV DMV membranes, the first time this has been observed in cells infected with a *GammaCoV*. This confirms that DMV pores are conserved features across the CoV family and is consistent with the recent identification of DMV pores in more distantly related arteriviruses^39^. Although insufficient pores were imaged for STA, initial analysis indicates they are very similar to *BetaCoV* DMV pores^24^. Further cryo-ET and STA will be required to allow a more detailed comparison of the pore structure. However, the lengths of the cytosolic prongs (10 nm) and the diameter of the IBV pore (20 nm) are comparable to those measured for other CoVs and the minimal DMV pore^16,24^. Furthermore, there were visible RNA filaments inside DMVs and RNA-like densities interacting with the luminal and cytosolic sides of the DMV pores as reported previously^11,17^. Therefore, the arrangement and structure of the *GammaCoV* DMV pore appear comparable to *BetaCoVs*.

In addition to the DMV pore, a range of DMV diameters were observed in both cryo-SXT and cryo-ET datasets. Although the mean DMV diameter was similar to reports from previous work^8,24,26,31^, a population of smaller DMVs was also identified, which was recapitulated in further RT ET samples. Significantly, these smaller DMVs contained a similar number of pores to the larger DMVs and this may give some important clues as to DMV biogenesis. It has been shown that *BetaCoV* nsp3 and nsp4 are sufficient to produce DMVs with the addition of nsp6 producing DMVs more similar to those produced during viral infection^16,25,27,29,30,38^. Furthermore, DMV biogenesis requires cleavage of the nsp3-nsp4 junction, which may facilitate bending of paired membranes into a DMV during pore formation^16,28,30^. As the large and small IBV DMVs contained comparable numbers of visible pores, this supports a model where DMV and pore formation are simultaneous. In addition to this, the small DMVs lacked obvious RNA filaments, and some instead contained a dense material. The identity of the dense material is unknown, but it is possible that it is either unprocessed polyprotein 1a/1ab or compacted viral RNA. Either way, this may indicate that smaller DMVs are initially formed, which subsequently mature and expand into larger DMVs^40^ perhaps as a result of pressure induced by the process of viral RNA synthesis within the lumen, as has been proposed for ROs from other +RNA viruses^41^. It is an interesting hypothesis that will require experimental testing in the future.

Using cutting edge cryo-bioimaging approaches we have been able to advance understanding of important and conserved features of the CoV replication cycle. Most significantly, we have described for the first time for any CoV a complex triple-layered protein lattice embedded within spherules. We also confirmed the presence of a double membrane spanning protein pore in the DMVs of a *GammaCoV*. Although this was perhaps expected, it is nonetheless a highly significant result. Further, we observed potential formation intermediates or immature DMVs at a high level of detail providing new leads to follow up into the process of DMV biogenesis and maturation. In summary, we have provided an enhanced level of structural understanding to underpin further functional characterisation of these conserved and intriguing elements of the CoV replication organelle.

## Methods

### Ethics statement

Use of chickens for production of primary cells was carried out in accordance with the Home Office guidelines of the United Kingdom (UK) and UK Animals (Scientific Procedures) Act 1986. Licensed facilities at the Pirbright Institute under establishment license number X24684464 were used for animal housing and all procedures. Embryonated hens’ eggs for primary cell production were purchased from The National Avian Research Facility (NARF) located at The Roslin Institute. Embryonated hens’ eggs for the propagation of viruses were purchased from VALO Biomedia, Germany.

IBV reverse genetics and modification of the IBV genome to include the meGFP tag have been risk assessed and approved by the Pirbright Institute’s Biological Agents and Genetic Modification Safety Committee (BAGMSC). In accordance with this risk assessment, virus work was carried out in containment level 2 facilities.

### Cells, embryonated eggs and viruses

Primary chick kidney (CK) cells were prepared from 2-3-week-old specific pathogen-free (SPF) Rhode Island Red (RIR) chickens by the Cell Culture Science Technology Platform at the Pirbright Institute^42^. Avian DF1 cells (LGC Standards Ltd., Teddington, UK) are a continuous cell line derived from chick embryo fibroblasts^43^. SPF embryonated hens’ eggs were ordered from VALO BioMedia GmbH and were set by the Animal Services Unit at the Pirbright Institute.

Recombinant IBV (rIBV) Beau-R (Accession: AJ311317.1)^22^ is referred to as wild type (WT) in this study. Beau-R expressing meGFP tagged nsp2 (Beau-R-eGFP-nsp2) was generated using a reverse genetics system based on a recombinant vaccinia virus (rVV) containing a complementary DNA (cDNA) copy of the complete IBV genome, as described previously^22,44,45^. The meGFP open reading frame (encoding the K206A mutation) followed by a 3x alanine linker (GCAGCTGCC) was inserted at nucleotide position 531 of the Beau-R sequence producing an N-terminal in-frame fusion with nsp2 (Figure 1). Stocks of the rIBVs were made in CK cells and embryonated eggs and titrated by plaque assay in CK cells, as described previously^46,47^.

### Growth kinetics

Confluent CK or DF1 cells in 6 well plates were infected with virus at 1 x 10^5^ PFU (plaque forming units) giving an approximate MOI of 0.1. Cell supernatants were harvested at specified timepoints, and the presence of progeny virus was assayed by plaque assay on CK cells. For RT-qPCR, RNA was extracted (Qiagen) and reverse transcribed (Super Script IV), according to the manufacturer’s instructions, followed by qPCR using primers and probe specific for the IBV 5ʹ UTR, as described previously^8,48^. Sample Ct values were interpolated from a standard curve to measure the copy number.

### Immunofluorescence imaging

Cells seeded onto coverslips were infected. At the indicated timepoint, cells were fixed with 4% paraformaldehyde and labelled with primary and secondary antibodies as indicated. For BrU labelling of nascent viral RNA, cells were treated with 4 mM bromouridine (BrU; Sigma Aldrich, 850187) and 15 µM actinomycin D (ActD; Sigma Aldrich, A9415) at 30 minutes prior to each timepoint. As described previously, for BrU detection, labelling steps were carried out in an RNAse free environment^13^. Cells were visualised using a Stellaris 5 confocal microscope (Leica Microsystems) and images were collected in the Leica LAS X software.

Alternatively, cells were imaged on a Leica SP8 gSTED 3X microscope with inverted stand. Images were collected in Leica LAS X software and then deconvolved using Huygens Professional software 18.10 (Scientific Volume Imaging).

Primary antibodies were used as follows: anti-BrU (Bromodeoxyuridine) (Sigma, 11170376001) at 1:50, anti-S 26.1 (Wageningen university and research) at 1:100, anti-N 48.4 (Wageningen university and research) at 1:1000, anti-M 25.1 (Wageningen university and research) at 1:1000, anti-E AF12 (Immunological Toolbox, Pirbright) at 1:50. Alexa Fluor conjugated secondary antibodies (Thermo Fisher Scientific) were used at 1:500.

### Correlative light and electron microscopy (CLEM)

DF1 cells were infected in dishes containing gridded coverslips (Mattek, P35G-1.5-14-C-GRD). At the specified time post infection, the nuclei were labelled with Hoechst 34580 (Thermo Fisher Scientific, H21486) at 5 μg/ml for 15 min and the media was replaced with Leibovitz’s L-15 medium without phenol red (Thermo Fisher Scientific, 21083027). Selected grid squares were imaged live using a Leica STELLARIS 5 laser scanning confocal microscope. Grid coordinates were captured with a low magnification brightfield image, followed by high magnification image acquisition of cells of interest within each selected grid square. At the specified times, samples were fixed with 2% glutaraldehyde and resin embedded (Agar Scientific, AGR1031) as described previously^18^. Sections (80 nm) were cut from appropriate grid squares, collected onto copper grids (Agar Scientific) and stained using a Leica EM AC20 instrument. The specific cells imaged by confocal microscopy were located and imaged using a Talos L120C TEM (Thermo Fisher Scientific) at 120 kV with a 4k × 4K Ceta CMOS camera, as described in^8^. Composite images were compiled using Adobe Photoshop.

### Room temperature electron tomography

CK cells were seeded onto 13 mm Thermanox coverslips (Thermo Fisher Scientific, UK) and infected with Beau-R. At 18 hpi, cells were fixed in phosphate buffered 2% glutaraldehyde and embedded in resin. Thick sections (250 nm) were cut and collected onto Formvar/carbon coated copper 2×1mm slot grids (EMS) then 10 nm gold antibody (Aurion) added to the grids to act as fiducial markers. Grids were imaged in a JEOL 2100F TEM at 200 kV with a TVIPS F416 digital camera. Data sets were collected using Serial EM, with acquisitions every 1° over a 120° tilt series. Tomogram reconstruction and DMV measurements were carried out using IMOD. DMVs were measured using line contours throughout the tomogram, with the longest contour being recorded as the diameter. Line contours were only drawn for DMVs whose widest diameter was within the tomogram depth. Data was imported into Graphpad Prism and violin plots produced.

### Cryo-soft X-ray tomography

DF1 cells (10^5^ cells) were seeded onto R2/2 Au200F1 grids (Quantifoil) in 6 well plates and infected with Beau-R-eGFP-nsp2. Cells were vitrified on a Leica GP2 at -170 °C with a blot time of 0.5-2s and addition of 250 nm gold fiducials (BBI Solutions, EM.GC250). Grids were screened for eGFP signal under the cryo-SIM developed in-house at beamline B24 at Diamond Light Source as described previously^49,50^. The fluorescently screened grids were imaged on the UltraXRM-S/L220c X-ray microscope (Carl Zeiss X-ray Microscopy) at beamline B24 set up with the 25 nm zone plate. Samples were illuminated with 500 eV X-rays (λ = 2.48 nm) for 0.5 or 1 s per projection. Transmitted images were collected using a 1024B Pixis CCD camera (Princeton instruments) at 10 nm/pixel. Tilt series were collected in increments of 0.2° or 0.5° per image, with tilt angles of ±60°. Reconstruction of the X-ray tilt series into tomograms was completed using eTomo in IMOD (version 4.9.2; ^51^). Tomograms were visualised in ImageJ^52^ and segmentations were completed using 3dmod in IMOD.

### Cryo-FIB lamella preparation

DF1 cells (10^5^ cells) were seeded onto R2/2 Au200F1, Au300, or Au200 grids (Quantifoil) in 6 well plates and infected with Beau-R-eGFP-nsp2. Cells were vitrified on a Leica GP2 at - 170 °C for with a blot time of 8-10 s and addition of 5 µl of 10 % glycerol (Merck, G2025) in PBS containing a 1:10 dilution of Dynabead myOne 1 µm diameter beads (Thermo Fisher Scientific, 65001). Grids were screened for eGFP fluorescence using a Leica EM Cryo-CLEM. Grids were then loaded into the Scios DualBeam cryo-FIB-SEM system (Thermo Scientific). Grids were sputter coated with a layer of platinum (0.1 mbar Ar, 10 mA, 500 V, for between 30 and 60 s) before loading onto the Scios cryo stage. Before milling, a layer of organoplatinum precursor was deposited on the grid for 5-10s using the in-chamber gas injection system. The accessible cells of interest were found using the SEM (operated at between 2 and 5 kV, and 13 and 25 pA) and 30 keV FIB images were collected at a milling angle of 12°. Leica THUNDER Imager EM cryo-CLEM z-stacks were correlated to the FIB view images using 3D Correlative Toolbox (3DCT, ^53^). Milling regions were accurately placed based on the 3D correlation to the fluorescence microscopy. Milling was performed using the Ga+ ion beam, operated at 30 kV. Milling was performed in steps, with ion beam currents selected according to the separation between milling regions (Table 1).

**Table 1:**
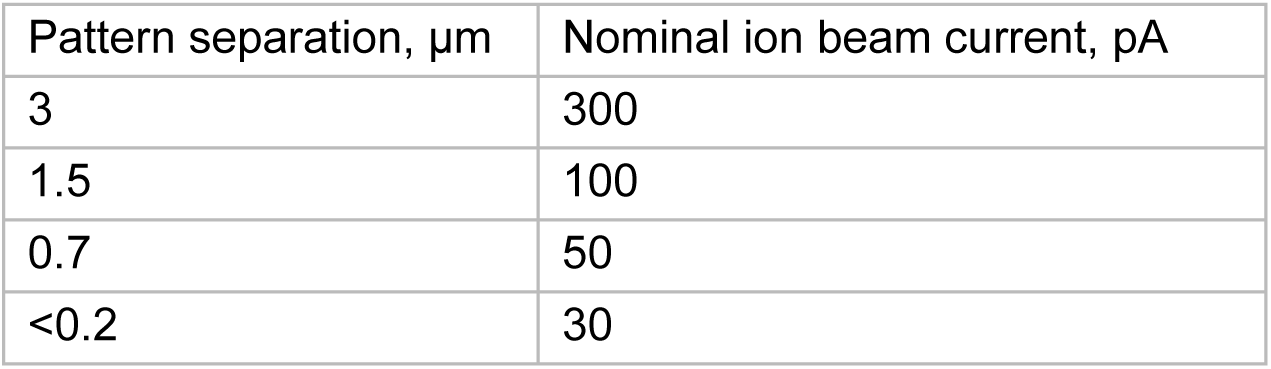
Sequential ion beam currents used when producing lamellae.

### Cryo-electron tomography data acquisition

Grids with lamellae were loaded into an autoloader cassette so that the milling direction was perpendicular to the microscope tilt axis to give the maximum tilt range during data collection. A Titan Krios electron microscope (Thermo Fisher Scientific) located at eBIC (Krios III) was used to collect tilt series at 300 keV with a selectris-X filter and falcon 4i detector. Samples were pre-tilted to 12° to compensate for the milling angle. Tilt series were collected at a pixel size of 3.05 Å (nominal magnification of 42kx), with a defocus range of -5 to -2 µm, a tilt range of ±50° with 2° increment, and an electron dose of 2.06 e-/Å2/tilt.

### Cryo-electron tomography image processing

On-the-fly processing of tomograms was done using the eBIC Murfey pipeline^54^ to assess data collection quality. Tilt series were motion corrected using MotionCor2 ^55^, CTFFind4^56^ was used for CTF estimation, and AreTomo^57^ was used for alignment and reconstruction via patch tracking. Automatic denoising is performed with Topaz^58^ using a pre-trained model. The pipeline also uses IMOD^59^ for basic micrograph manipulations.

For membrane segmentations, reconstructed tomograms were also denoised using Isonet^60^. The membranes were then segmented using Membrain-seg^61^. Other segmentations were also completed using 3dmod in IMOD (version 4.9.2;^51^). UCSF ChimeraX^62^ was used to model reconstructed tomograms, membrane segmentations, subvolume averages, and AlphaFold and PDB structures.

### Subtomogram averaging

For subtomogram averaging (STA), RELION-5^63^ was used. In this pipeline, MotionCor2^55^, CTFFind4^56^, and AreTomo^57^ were also used for reconstruction and cryoCARE0.3.0^64^ was used for denoising. Particle picking was done using a Napari-based viewer^65^ and was completed within the pipeline and STA was completed on the extracted subvolumes as described for RELION-4^66^. Spherule structures were picked using the oversampling around a sphere technique as described in previous work^67,68^. The centre of each spherule was marked and the radius was set at the inner spherule membrane. Particle spacing was set at 40 Å for the extraction of subtomograms around the spheres. The spherical picking also provided a prior angle for each of the particles so that the particles were orientated with their Z-axis normal to the surface of the pseudo-spheres. Using this technique, 98567 particles were picked and subtomograms were extracted at bin 4 (12.2 Å/px). From these subtomograms, a reconstruction was made to act as a reference map for 3D classification.

The best class was chosen as a new reference map for further 3D classification. Once more, the best class was used as an updated reference map for 3D classification. The best class contained 13966 particles and from this refined optimisation set, particles were extracted at bin 2 (6.1 Å/px, Fig. s2). Following particle reconstruction at bin 2, the reconstruction was used as a reference map for 3D auto-refinement. Next, duplicate particles with a minimum inter-particle distance of 40 Å were removed resulting in 3796 particles for ultimate 3D reconstruction at bin 2, masking and refinement. The final resolution of the refined map was 23 angstroms.

## Acknowledgements

We would like to thank the Pirbright Institute Cell Culture Science Technology Platform (STP) for weekly preparation of primary chick kidney cells and other cell culture support and the Bioimaging STP for imaging advice and support. We would like to thank the Coronavirus Cellular Biology and Coronavirus research groups for useful discussions. We acknowledge Diamond for access and support of the cryo-EM facilities at the UK national electron Bio-Imaging Centre (eBIC), proposal NT26538.

## CRediT author contributions

**Conceptualisation**: HJM, DC, MH, JMG, PCH, PMP. **Investigation**: PMP, JW, NM, ND, SK. **Methodology**: PMP, JW, ND, SK, PM, JBG, MH, DC, HJM. **Formal analysis**: PMP, JW, NM, ND, MH, DC, HJM. **Visualisation**: PMP, JW, NM, HJM. **Supervision**: HJM, DC, MH, JMG, PCH. **Project administration**: HJM, DC, MH. **Resources**: HJM, DC, MK.

**Funding acquisition**: HJM, DC. **Writing – original**: PMP, HJM. **Writing – review and editing**: PMP, JW, NM, ND, SK, PM, JBG, PCH, JMG, MH, DC, HJM.

---

**Supplementary Figure 1:**
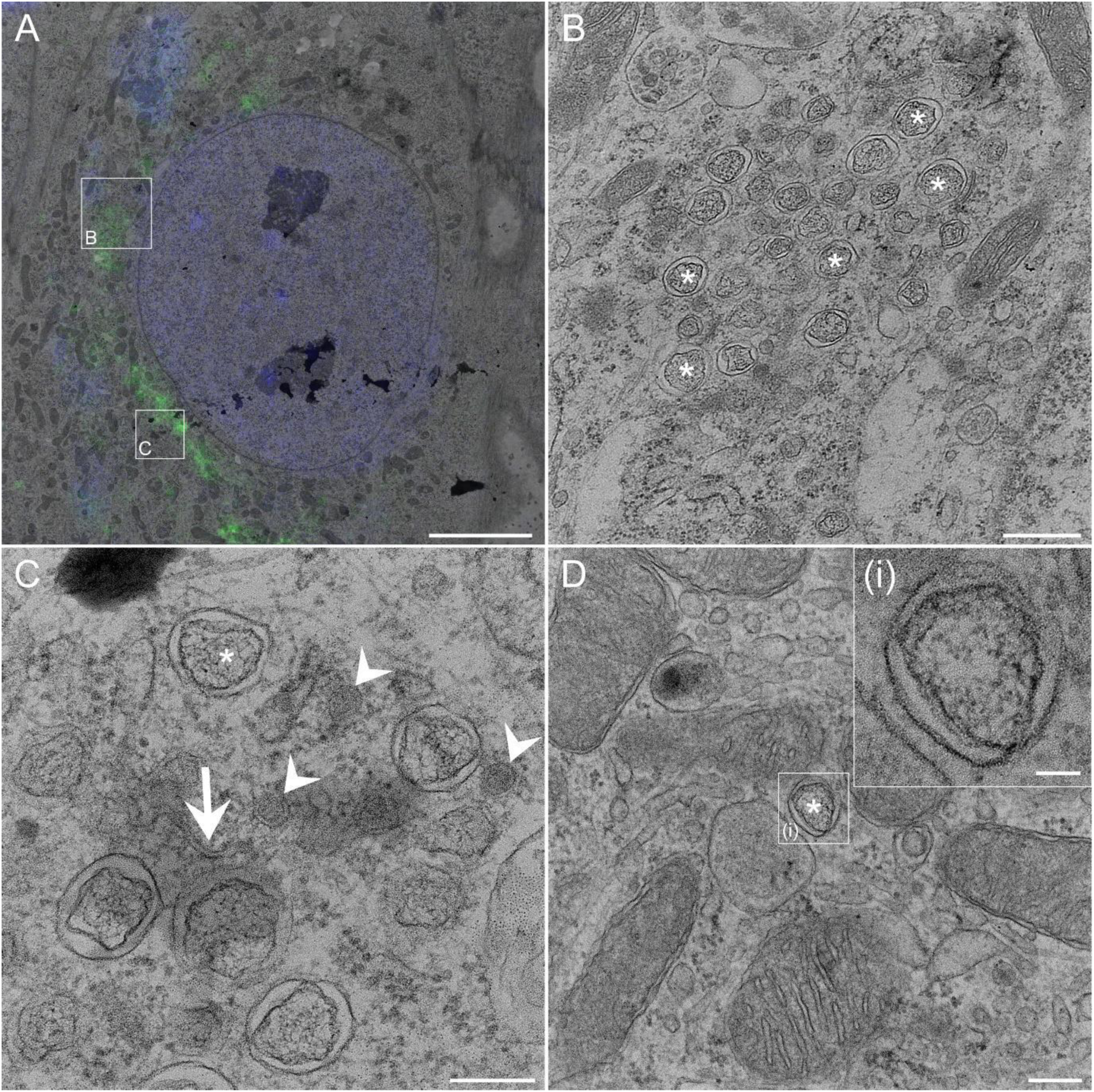
IBV ROs can be detected from 4 hours post infection. CK cells were infected with Beau-R-eGFP-nsp2 (green), nuclei were labelled with Hoechst (blue) and live cells were imaged. Cells were then fixed and resin embedded at 6 h pi (**a**-**c**). (**a**) Fluorescence images were overlaid onto electron micrographs to localise replication organelle membranes. Boxed areas were imaged at higher magnification (**b**, **c**). (**d**) CKCs were infected with Beau-R-eGFP-nsp2 and fixed and resin embedded at 4 hpi, highlighting a small DMV (**di**). Asterisks indicate DMVs, white arrows indicate zER and white arrowheads indicate spherules. Scale bar indicates (**a**) 1 µm, (**b**, **d**) 500 nm, (**c**) 200 nm, (**di**) 50 nm.

**Supplementary Figure 2:**
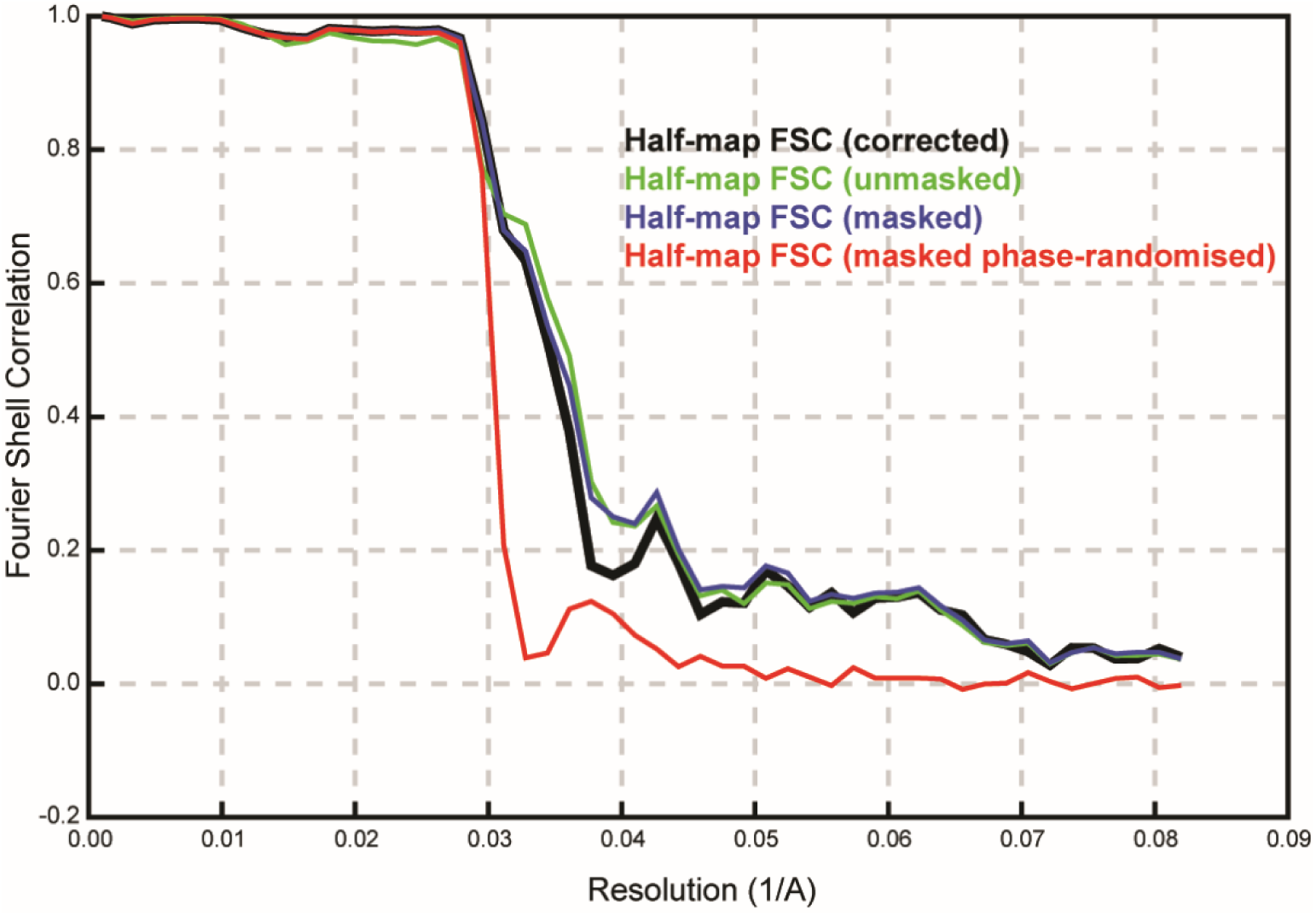
Half-map FSC curve for the STA map of the spherule double membrane. FSC curves are given for map resolution (half-map FSC) and the resolution values are coloured according to the key.

**Supplementary Figure 3:**
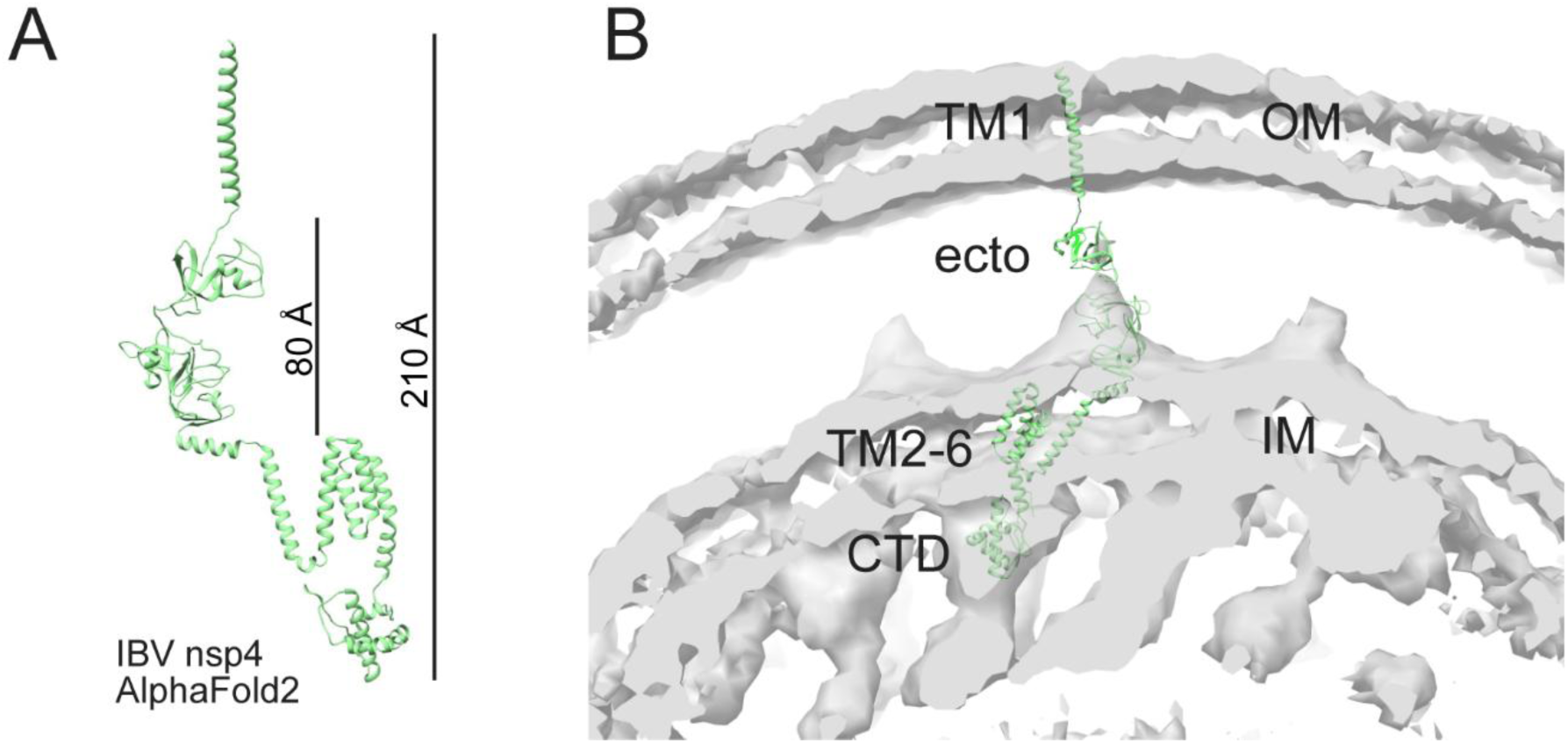
Hypothesised model of IBV nsp4 fit into the spherule triple-layer lattice density map. (**a**) AlphaFold2 predicted structure for IBV nsp4 with extended orientation of transmembrane domain 1. (**b**) An estimated fit model of IBV nsp4 into the spherule STA density model with TM1 in the outer membrane (OM), the ectodomain (ecto) in the intermembrane space, TM2-6 in the inner membrane (IM), and the C-terminal domain (CTD) in one part of the z-layer electron density under the inner membrane.

